# Enhancer evolution as a driving force for lineage-specific paralog usage in the central nervous system

**DOI:** 10.1101/2023.04.20.537653

**Authors:** Chika Fujimori, Kohei Sugimoto, Mio Ishida, Christopher Yang, Daichi Kayo, Soma Tomihara, Kaori Sano, Yasuhisa Akazome, Yoshitaka Oka, Shinji Kanda

## Abstract

Expression patterns of paralogous genes in the functionally homologous cells sometimes show differences across species. However, no reasonable explanation for the mechanism underlying such phenomena has been discovered. To understand this mechanism, the present study focused on the hypophysiotropic GnRH neurons in vertebrates as a model. These neurons express either *gnrh1* or *gnrh3* paralogs depending on species, and apparent switching of the expressed paralogs in them occurred at least four times in vertebrate evolution. First, we found redundant expressions of *gnrh1* and *gnrh3* in a single neuron in piranha and hypothesized that this situation may indicate an ancestral condition. We tested this hypothesis by examining the activity of piranha *gnrh1/gnrh3* enhancers in zebrafish and medaka, in which the two *gnrh* paralogs are not co-expressed. Here, the *gnrh1/gnrh3* enhancer of piranha induced reporter RFP/GFP co-expressions in a single hypophysiotropic GnRH neuron in both zebrafish and medaka. From these results, we propose that long-lasting (∼550 My) redundancy after *gnrh1/3* duplication in 1R/2R WGD may be the key to apparent switching of the paralog usage among the present-day species. Moreover, interspecies analyses of enhancers indicated that the loss of enhancers rather than changes in trans-regulatory elements drove the role-division of these paralogs.

## Introduction

The theory of evolution by gene duplication tells us that, after the gene duplication, which is a main source of novel genes, one of the duplicated genes undergo degradation (non-functionalization) or acquisition of novel function (neo-functionalization), while the other retains the original function or both genes divide up the original function/expression pattern (sub-functionalization) [1-3]. In any case, duplicate genes are supposed to lose their functional redundancy during evolution by accumulating mutations over a relatively short period of time [3-5], and once their fates have been fixed, their functional roles cannot be changed or swapped. Contrary to this principle, apparently strange phenomenon is observed in a peptide hormone called gonadotropin-releasing hormone (GnRH), which is a key molecule for reproduction. In all vertebrate species that have been investigated thus far, one of the *gnrh* paralogous genes is expressed in GnRH neurons in hypothalamus/preoptic area (POA), which project to the pituitary and control gonadal functions by inducing the release of gonadotropins (a so-called hypophysiotropic function). Here, the functional subtype (paralog) of the *gnrh* gene for pituitary regulation has been shown to vary across species, and has apparently switched several times during vertebrate evolution [6, 7]. There are three *gnrh* paralogs (*gnrh1/2/3*) which appeared after the first/second-round (1R/2R) whole genome duplication (WGD) ∼550 million years ago (Mya) [8-11] (Fig. S1). In many species including tetrapods and some teleosts, *gnrh1* is expressed in the hypophysiotropic GnRH neurons as the main regulator of gonadotropin release from the pituitary [12-17]. However, some teleost species including zebrafish express *gnrh3* in hypophysiotropic GnRH neurons instead of *gnrh1*. Interestingly, all species that express *gnrh3* in hypophysiotropic GnRH neurons have genetically lost *gnrh1* [18-20] (Fig. S1). Since the peptide sequences among different GnRH paralogs are highly conserved, and their ligand-receptor relationship has been considered promiscuous (all the GnRH subtypes basically show similar actions on the GnRH receptors) [21, 22], *gnrh3* has been considered to have compensated for the loss of *gnrh1* during evolution [6, 7]. However, no study has yet provided a reasonable explanation as to why functionally homologous neurons express different paralogs among different species.

Such a seeming “switch” in the usage of paralogs in the hypophysiotropic GnRH neurons has occurred at least four times in the evolutionary lineage of teleosts (Fig. S1). If we can find a species whose hypophysiotropic GnRH neurons co-express both *gnrh1* and *gnrh3*, it should give strong supporting evidence for explaining the mechanism for this seeming switch of paralog usage among different species, e.g., co-expression of *gnrh1* and *gnrh3* in the hypophysiotropic GnRH neurons in the common ancestor may play a permissive role in each species to lose one of them. Interestingly, we found reports in two characiform species (order Characiformes); in pacu (*Piaractus mesopotamicus*, family Serrasalmidae) three different forms of GnRH peptides were identified by using HPLC analysis followed by mass spectrometry [23], whereas *gnrh1* gene could not be identified in another Characiform family, Characidae species, including *Astyanax altiparanae* [24] and Mexican tetra, in which *gnrh1* is not found in Ensembl genome database (Fig. S1). Since this different pattern of paralog usage occurred in different species in the Characiformes, we surmised that species in the family of Serrasalmidae, as presumed ancestral type, may provide a hint toward understanding the mechanism underlying switches in *gnrh* paralog usage.

In the present study, to understand the evolutionary mechanism of such switches in paralogous-gene usage, we analyzed *gnrh1* and *gnrh3* system of a Serrasalmidae fish, the red-bellied piranha (*Pygocentrus nattereri*). First, we found in piranha that *gnrh1* and *gnrh3* are co-expressed in the hypophysiotropic GnRH neurons, which is suggestive of a possible existence of common ancestral teleosts having the similar situation. We surmised that this finding of the co-expression of two paralogs may explain the seeming switch in *gnrh* paralog usage in the hypophysiotropic neurons, because this situation may play a permissive role in the loss of either *gnrh1* or *gnrh3* expression/gene for regulating gonadotropin release without severe functional defects. Furthermore, to find out experimental support for our surmise above, the enhancer activity of the piranha *gnrh* gene was tested in other fishes, medaka and zebrafish, which exhibit different *gnrh1/3* systems.

## Results

### In piranha, gnrh1 and gnrh3 are co-expressed in the POA GnRH neurons

Sequences of piranha *gnrh1* and *gnrh3* mRNA were successfully determined by 5’ and 3’ RACE method (Fig. S2A, B). Also, partial sequences of head-and-tail-light tetra (*Hemigrammus ocellifer*, Characidae) *gnrh3* mRNA were isolated by 3’ RACE followed by RT-PCR (Fig. S2C). Based on the precursor sequences, the analyses of alignment (Fig. S2D, E) and phylogenetic tree (Fig. S2F) strongly suggests that these genes are piranha *gnrh1*, piranha *gnrh3* and head-and-tail-light tetra *gnrh3. In situ* hybridization using piranha *gnrh1* and *gnrh3* specific probes demonstrated that both *gnrh1* and *gnrh3* mRNA-expressing neurons are localized in the terminal nerve (TN) as well as the POA in the piranha brain (Fig. 1A, B). Since some *gnrh1*-expressing and *gnrh3*-expressing neurons appeared to be localized in adjacent areas of the brain, we performed double *in situ* hybridization of *gnrh1* and *gnrh3* to examine whether they are co-expressed in the same neuron. Double *in situ* hybridization clearly demonstrated that some neurons in both TN and POA co-expressed *gnrh1* and *gnrh3* mRNA (Fig. 1C, D, yellow arrows). It should be noted that large neurons of the TN only expressed *gnrh3* mRNA, whereas smaller cells in both the TN and the POA co-expressed *gnrh1* and *gnrh3* mRNA.

**Fig. 1.**
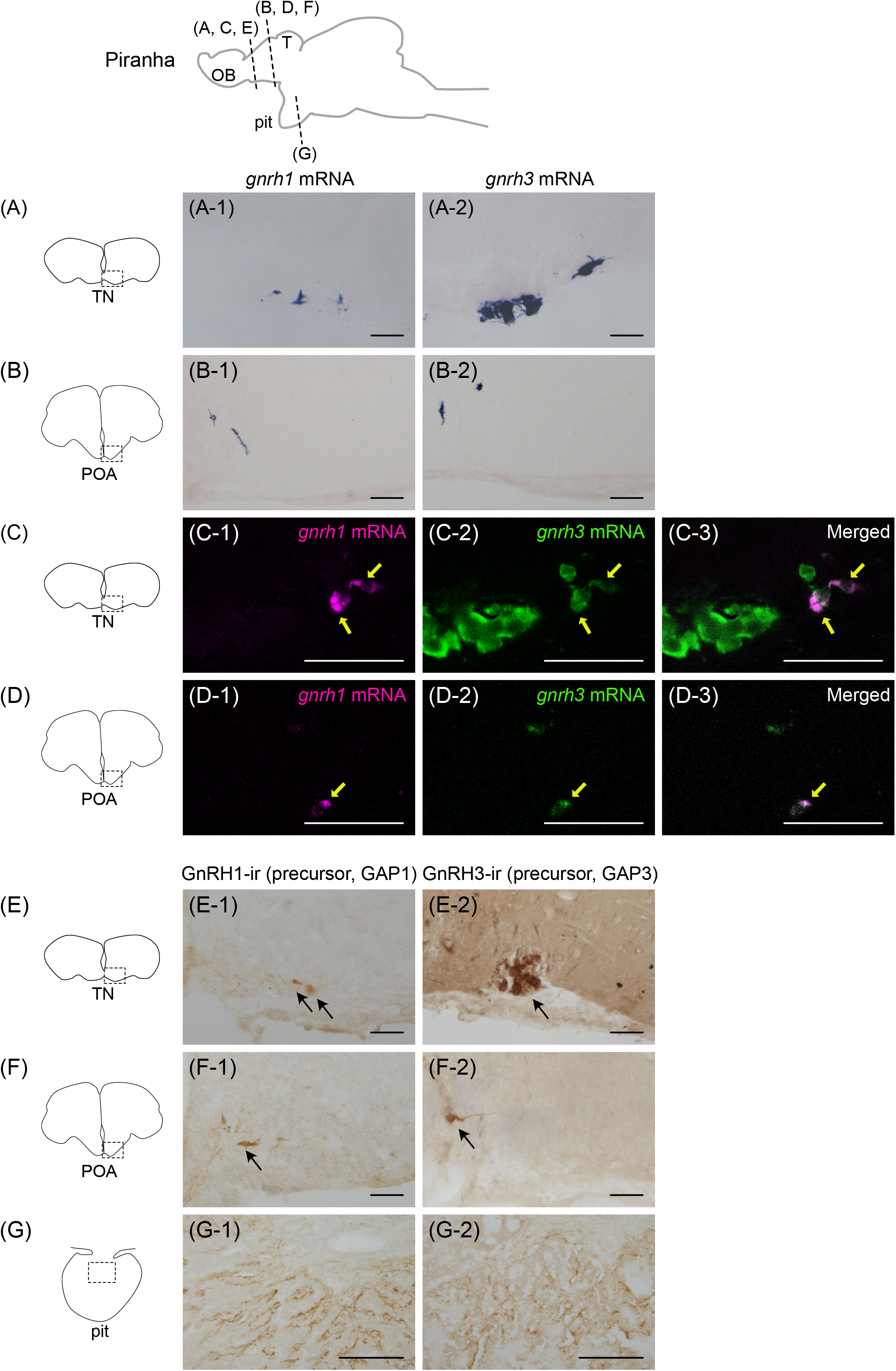
*Gnrh1* and *gnrh3* are co-expressed in the same hypophysiotropic neurons in piranha. **(A, B)** Both *gnrh1* **(A-1, B-1)** and *gnrh3* **(A-2, B-2)** mRNA are expressed in the terminal nerve (TN) **(A)** and the preoptic area (POA) **(B). (C, D)** Double *in situ* hybridization indicates that *gnrh1* and *gnrh3* mRNA are co-expressed in the same neuron in the TN (**C-1**, *gnrh1* mRNA; **C-2**, *gnrh3* mRNA; **C-3**, merged) and the POA (**D-1**, *gnrh1* mRNA; **D-2**, *gnrh3* mRNA; **D-3**, merged). Yellow arrows indicate the cells that co-express *gnrh1* and *gnrh3* mRNA. Note that neuronal cluster with giant cell bodies only express *gnrh3*. **(E, F)** Immunohistochemistry using precursor of GnRH1 **(E-1, F-1)** or GnRH3 **(E-2, F-2)**-specific antibody labeled the cell bodies in the TN **(E)** and the POA **(F)**, which is consistent with the localization indicated by *in situ* hybridization. Arrows indicate cell bodies. **(G)** Both GnRH1-**(G-1)** and GnRH3-**(G-2)** immunoreactive (ir) fibers are observed in the pituitary, which suggests the redundant regulation of LH release by GnRH1 and GnRH3 peptides. Scale bars represent 100 μm **(A, B, E, F, G)** and 50 μm **(C, D)**, respectively. OB, olfactory bulb; T, telencephalon; pit, pituitary.

### Both GnRH1-immunoreactive(ir) and GnRH3-ir neuronal fibers project to the pituitary in piranha

By immunohistochemistry using the newly generated antibodies against piranha GnRH1 or GnRH3 precursor (GAP1/GAP3), we analyzed the axonal projection of GnRH1 and GnRH3 neurons in piranha after scrutinizing the specificities of these antibodies by pre-absorption with the peptides (Fig. S3). Both the GnRH1-ir and GnRH3-ir cell bodies were localized in TN as well as POA (Fig. 1E and F), and the densely labeled axonal fibers of both GnRH1-ir and GnRH3-ir neurons were observed in the pituitary (Fig. 1G).

### Gnrh1 knockout piranha shows a similar pattern of GnRH innervation to that of head-and-tail-light tetra

Next, to investigate the functional compensation for the loss of the *gnrh1* by *gnrh3* gene gene, which supposedly occurred in Characidae linages (Fig. S1), we newly established an artificial fertilization method and generated *gnrh1* knockout (KO) piranha using CRISPR/Cas9. After microinjection of CRISPR/Cas9 mixture, F0 embryos were raised and incrossed to generate the F1 generation. In the F1 generation, PCR and subsequent sequencing analysis indicated that there were individuals that had frameshift mutation and/or result in non-functional GnRH peptide in the *gnrh1* gene (Fig. S2G), which we expected would fail to produce functional GnRH1 peptide. Immunohistochemistry for piranha GAP1 showed that neither GnRH1-ir cell bodies in the POA nor fibers in the pituitary were observed in *gnrh1*^*-/-*^ piranha (Fig. 2A-1, B-1), whereas the both were observed in *gnrh1*^*+/+*^ piranha(Fig. 2A-2, B-2). In both *gnrh1*^*-/-*^ and *gnrh1*^*+/+*^ piranha, GnRH3-ir fibers were similarly observed in the pituitary (Fig. 2C-1, C-2). The situation in this *gnrh1* KO piranha, which possess only *gnrh3* in hypophysiotropic GnRH neurons, was similar to the head-and-tail-light tetra, a closely related species that does not have *gnrh1* gene. Actually, in the POA of head-and-tail-light tetra, *gnrh3* was detected in the cell bodies of POA by *in situ* hybridization (Fig. 2D-1) as well as immunohistochemistry (Fig. 2D-2), and GnRH3-ir fibers were observed in the pituitary (Fig. 2E).

**Fig. 2.**
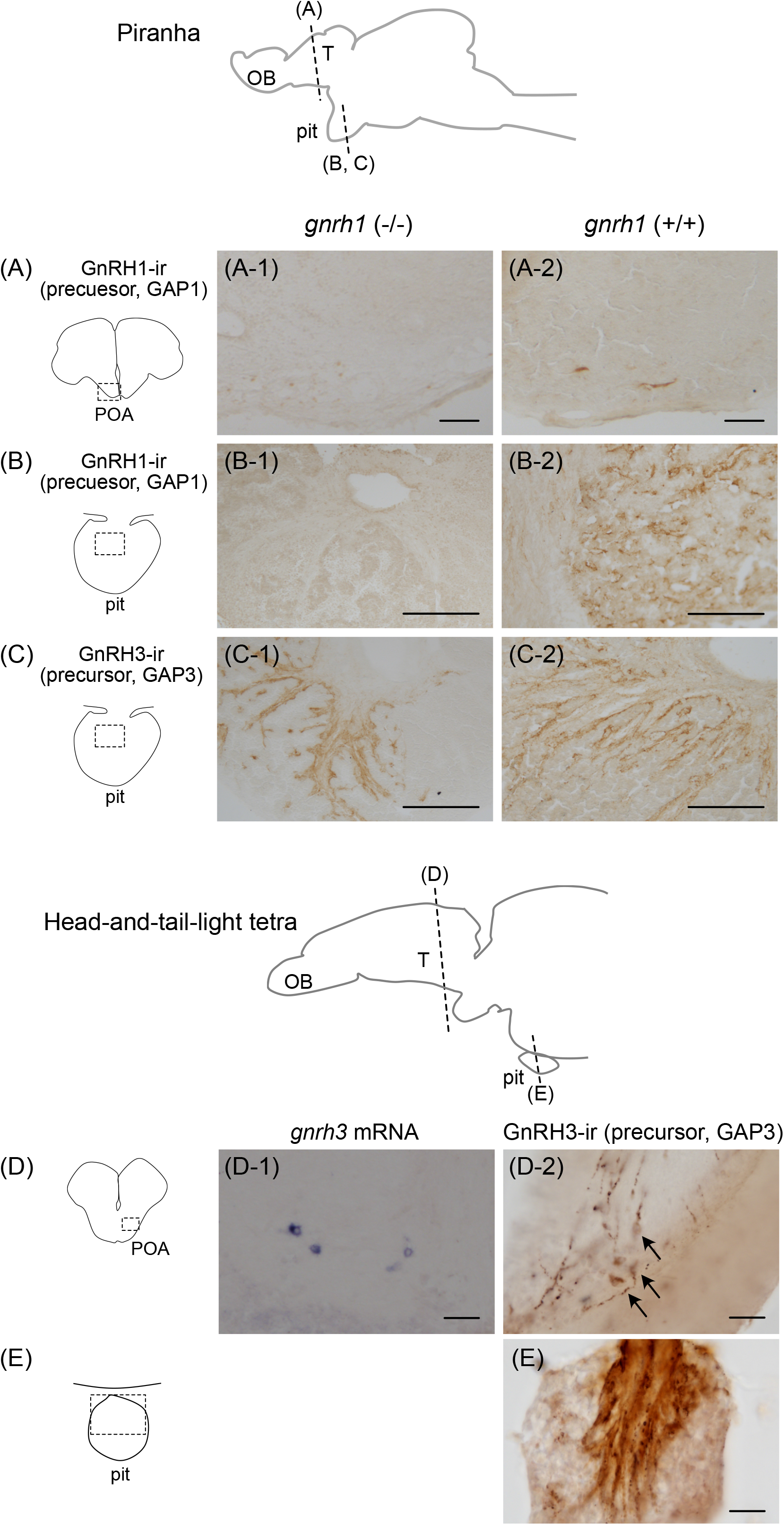
*Gnrh1* knockout (KO) piranha show innervation of GnRH3-ir fibers in the pituitary, similar to other characiform fishes. **(A)** GnRH1 precursor-ir cell bodies are not observed in the *gnrh1* KO piranha **(A-1)**, while they are observed in the wild type **(A-2). (B)** GnRH1 precursor-ir fibers are not found in the pituitary of the *gnrh1* KO **(B-1)**, while they are observed in the wild type **(B-2). (C)** GnRH3-ir fibers are observed in the pituitary of both *gnrh1* KO **(C-1)** and wild type **(C-2)** piranha. **(D)** In head-and-tail-light tetra, *in situ* hybridization **(D-1)** and immunohistochemistry **(D-2)** shows that *gnrh3/*GnRH3-expressing neurons are localized in the POA. **(E)** Immunohistochemistry indicates that GnRH3 precursor-ir axonal projection is observed in the pituitary. Arrows indicate cell bodies. Scale bars represent 100 μm **(A-C)** and 20 μm **(D, E)**, respectively. OB, olfactory bulb; T, telencephalon; pit, pituitary.

### In zebrafish, piranha gnrh1 and ghrh3 enhancers can be activated in the POA GnRH neurons expressing gnrh3 mRNA

The fact that *gnrh1* and *gnrh3* were co-expressed in the hypophysiotropic GnRH neurons in piranha strongly suggests that the common ancestor of Ostariophysi possessed hypophysiotropic GnRH neurons expressing both *gnrh1* and *gnrh3*. From such ancestor, the GnRH system in the present ostariophysian fishes, in which either *gnrh1* or *gnrh3* are expressed in the hypophysiotropic GnRH neurons, is considered to have arisen as a consequence of a simple loss of either one of the two *gnrh* paralogs. We then examined whether the trans-regulatory elements in zebrafish hypophysiotropic GnRH neurons can still activate the anciently lost *gnrh1* enhancer to know whether this loss of cis-regulatory elements and coding region was the only cause of the loss of expression of *gnrh1*. We generated transgenic (Tg) zebrafish harboring RFP (dTomato) or GFP (EGFP) as reporter gene under the regulation of piranha *gnrh1* or *gnrh3* 5’ flanking region, Tg (*pngnrh1*:RFP) and Tg (*pngnrh3*:GFP), respectively (Fig. 3A). In the established transgenic zebrafish, both piranha *gnrh1* (RFP) and *gnrh3* (GFP) transcriptional activities were observed in the cell bodies of endogenous GnRH3 neurons in POA (Fig. 3B, C), indicating that piranha *gnrh1* and *gnrh3* enhancers were activated in zebrafish GnRH neurons. These results strongly suggest that trans-regulatory elements activating both *gnrh1* and *gnrh3* enhancers are conserved even though the *gnrh1* gene has already been lost in zebrafish. Furthermore, in the double transgenic zebrafish, Tg (*pngnrh1*:RFP; *pngnrh3*:GFP), RFP and GFP expression was co-localized in both cell bodies in the POA (Fig. 3D) and the axons in the pituitary (Fig. 3E).

**Fig. 3.**
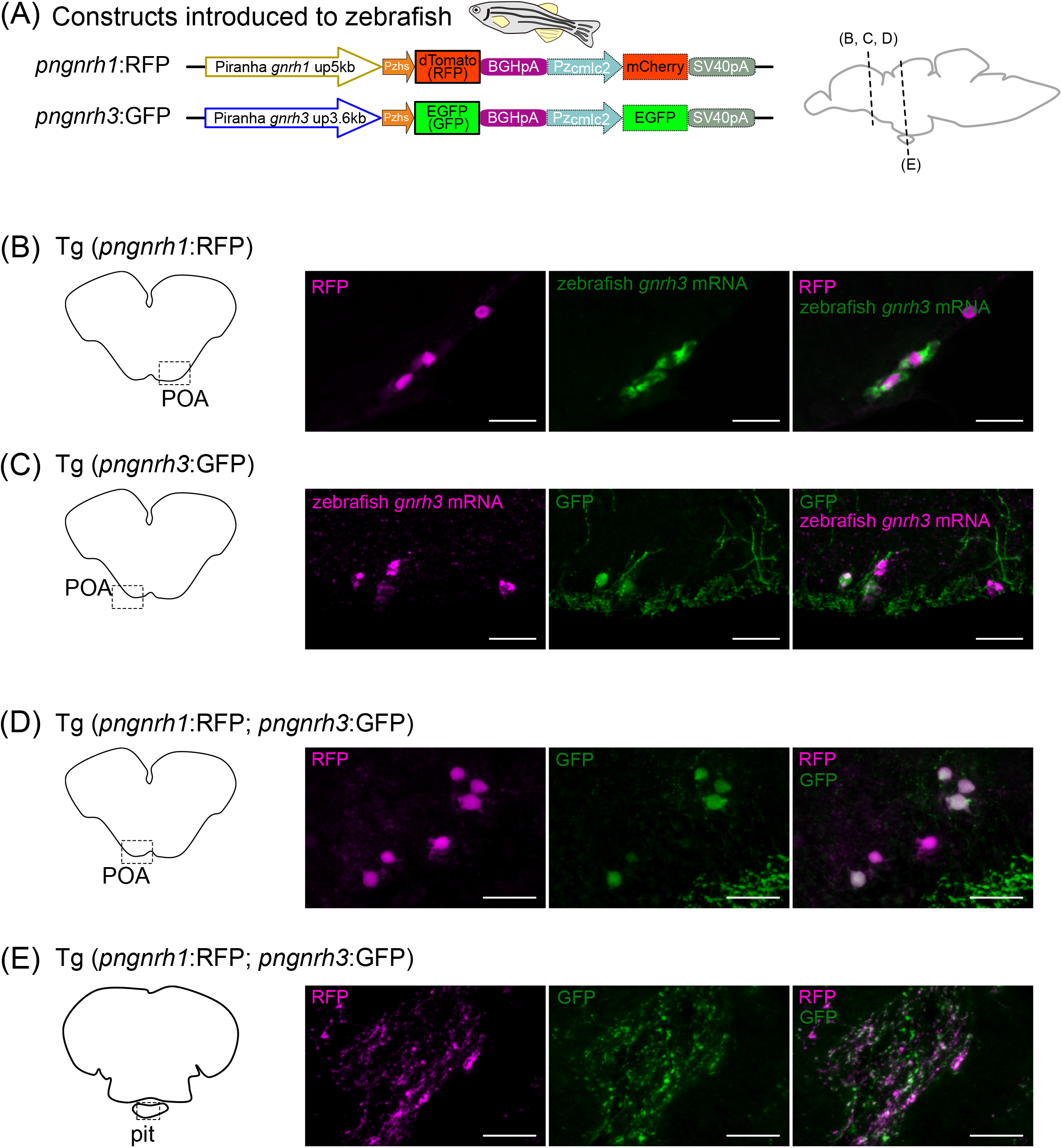
Examination of enhancer activity of *gnrh1* and *gnrh3* of piranha (*pngnrh1*/*pngnrh3*) in zebrafish. **(A)** The constructs used to generate transgenic zebrafish. Both constructs examine the enhancer activity of piranha *gnrh1* or *gnrh3* 5’flanking region by using basal promoter (zebrafish heat shock promoter, Pzhs) and a fluorescent protein (RFP/dTomato or GFP/EGFP). For screening of embryos, cardiac myosin light chain 2 promoter of zebrafish (Pzcmlc2), mCherry or EGFP and SV40 poly(A) signal were inserted downstream of the reporter construct. **(B, C)** Double labeling of piranha enhancer-induced fluorescent proteins and the intrinsic mRNA of zebrafish. **(B)** In Tg (*pngnrh1*:RFP) zebrafish, *pngnrh1* enhancer-induced RFP expression is observed in the GnRH3 neurons (*gnrh3* mRNA-expressing neurons) in the POA. **(C)** In Tg (*pngnrh3*:GFP) zebrafish, *pngnrh3* enhancer-induced GFP expression is observed in the GnRH3 neurons in the POA. **(D, E)** Analysis of the double transgenic zebrafish, Tg (*pngnrh1*:RFP; *pngnrh3*:GFP). **(D)** In Tg (*pngnrh1*:RFP; *pngnrh3*:GFP) zebrafish, some of the neurons in the POA express both RFP and GFP, suggesting that *pngnrh1* and *pngnrh3* enhancers are active in the same neurons. **(E)** In the pituitary of Tg (*pngnrh1*:RFP; *pngnrh3*:GFP) zebrafish, neuronal fibers that are labeled by both RFP and GFP are observed, suggesting that the RFP and GFP co-expressing neurons in the POA are hypophysiotropic. Scale bars, 20 μm.

### In medaka, both piranha gnrh1 and gnrh3 enhancers can be activated in the POA GnRH neurons expressing intrinsic gnrh1 mRNA

Although both *gnrh1* and *gnrh3* genes are retained in all acanthopterygian fishes examined thus far including medaka, hypophysiotropic GnRH neurons express *gnrh1* but not *gnrh3* in principle (Fig. S1) [12, 13, 15, 17]. To examine whether the lack of expression of *gnrh3* in the hypophysiotropic GnRH neurons in Acanthopterygii was due to changes in the enhancer sequence, we generated transgenic medaka, Tg (*pngnrh1*:RFP) and Tg (*pngnrh3*:GFP) and examined the enhancer activity of hypothetical ancient type piranha *gnrh1* and *gnrh3* enhancers in the hypophysiotropic neurons of medaka (Fig. 4A). In transgenic medaka, both piranha *gnrh1* and *gnrh3* enhancers induced RFP (dTomato)/GFP (EGFP) reporter expression in medaka *gnrh1* mRNA-expressing cell bodies in the POA (Fig. 4B, C). Moreover, in the double transgenic medaka, Tg (*pngnrh1*:RFP; *pngnrh3*:GFP), RFP and GFP expression was co-localized in both cell bodies in the POA (Fig. 4D) and the axons in the pituitary (Fig. 4E).

**Fig. 4.**
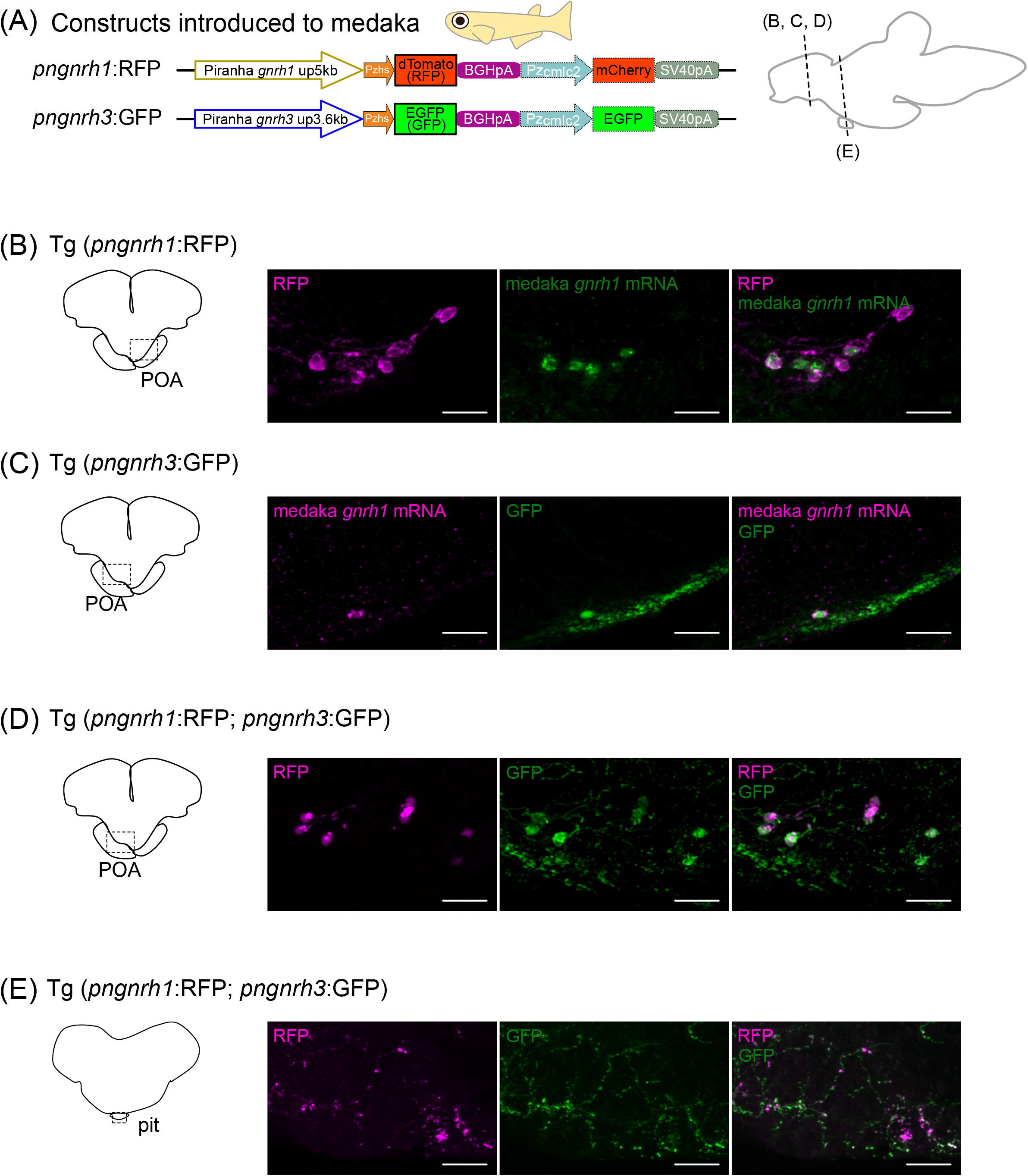
Examination of enhancer activity of *gnrh1* and *gnrh3* of piranha (*pngnrh1*/*pngnrh3*) in medaka. **(A)** The constructs used to generate transgenic medaka. Both constructs examine the enhancer activity of piranha *gnrh1* or *gnrh3* 5’flanking region by using a basal promoter (zebrafish heat shock promoter, Pzhs) and a fluorescent protein (RFP/dTomato or GFP/EGFP). For screening of embryos, cardiac myosin light chain 2 promoter of zebrafish (Pzcmlc2), mCherry or EGFP and SV40 poly(A) signal were inserted downstream of the reporter construct. **(B, C)** Double labeling of piranha enhancer-induced fluorescent proteins and the intrinsic mRNA of medaka. **(B)** In Tg (*pngnrh1*:RFP) medaka, *pngnrh1* enhancer-induced RFP expression was observed in the GnRH1 neurons (*gnrh1* mRNA-expressing neurons) in the POA. **(C)** In Tg (*pngnrh3*:GFP) medaka, *pngnrh3* enhancer-induced GFP expression was also observed in the GnRH1 neurons in the POA. **(D, E)** Analysis of the double transgenic medaka, Tg (*pngnrh1*:RFP; *pngnrh3*:GFP). **(D)** In Tg (*pngnrh1*:RFP; *pngnrh3*:GFP) medaka, some of the neurons in the POA expressed both RFP and GFP suggesting that *pngnrh1* and *pngnrh3* enhancers were active in the same neurons. **(E)** In the pituitary of Tg (*pngnrh1*:RFP; *pngnrh3*:GFP) medaka, neuronal fibers that are labeled by both RFP and GFP are observed, which suggests that the RFP and GFP co-expressing neurons in the POA were hypophysiotropic neurons. Scale bars, 20 μm.

## Discussion

Although many comparative anatomical studies have reported inconsistencies among species as to which paralog, *gnrh1* or *gnrh3*, is expressed in the hypophysiotropic GnRH neurons, the reason for this inconsistency has not been explained. In the present study, we found that both *gnrh1* and *gnrh3* are co-expressed in hypophysiotropic neurons in piranha. Given that the accidental *de novo* occurrence of co-expression of two specific genes in a single neuron should be extremely low, the co-expression of *gnrh1* and *gnrh3* in piranha should have been inherited since the gene duplication of *gnrh1* and *gnrh3* in the 1R/2R WGD. Furthermore, we analyzed the enhancer activities of piranha *gnrh1* and *gnrh3* in zebrafish, which have lost *gnrh1* gene during evolution, and medaka, in which *gnrh3* is not expressed in hypophysiotropic GnRH neurons. The results revealed that both piranha *gnrh1* and *gnrh3* enhancers can be activated in the hypophysiotropic GnRH neurons in medaka and zebrafish by their intrinsic trans-regulatory elements. This finding supports a hypothesis that the frequent switching of *gnrh* paralog usage in hypophysiotropic regulation during teleost evolution is due to the ancestral co-expression of *gnrh1* and *gnrh3* in hypophysiotropic neurons, which is still inherited by piranha. Thus, the analysis of these slowly evolving paralogous genes at the cellular level, including their expression as well as enhancer activities across species, is considered to provide a valuable model for understanding the mechanism of allocating distinctive roles to paralogs after gene duplication. The present approach should give us important insights into the evolutionary process of role-division of the paralog.

### Redundant expression of gnrh1 and gnrh3 in hypophysiotropic GnRH neurons in the hypothetical common ancestors may play a permissive role in seeming switch of GnRH paralog usage responsible for gonadotropin release

The present study demonstrated that piranha possess both *gnrh1* and *gnrh3* genes and show redundant expression of *gnrh1* and *gnrh3* genes in the hypophysiotropic GnRH neurons. Also, both GnRH1-ir and GnRH3-ir fibers densely projected to the pituitary (Fig. 1G), which are suggested to originate from POA *gnrh1* and *gnrh3* co-expressing neurons (Fig. 1D). On the other hand, closely related Characiform/Characidae species examined to date were suggested to have lost *gnrh1* according to genome database of Mexican tetra and RT-PCR results of neon tetra, head-and-tail-light tetra, and glow light tetra (Fig. S2H). Taken together, both *gnrh1* and *gnrh3* are considered to have been conserved from the time of emergence of Characiformes (125 Mya). Later, Serrasalmidae species (e.g. piranha and pacu) conserved them, whereas Characidae (e.g. neon tetra, Mexican tetra) lost *gnrh1* during their early evolutionary stages. Many other species so far examined in Ostariophysi lost either *gnrh3* (all species examined in Siluriform) or *gnrh1* (all species examined in Cypriniform) [20, 25-29], which indicates that they lost either *gnrh1* or *gnrh3* independently from the ancestor that co-express *gnrh1* and *gnrh3* in hypophysiotropic neurons like piranha observed in the present study. The existence of such dual-paralog co-expressing ancestors is likely to be the cause of the difference in the use of *gnrh* paralogs in hypophysiotropic GnRH neurons in the present-day species [6, 7, 30]. This hypothesis is supported by the expression analysis of *gnrh1* KO piranha and head-and-tail-light tetra in the present study. *Gnrh1* KO piranha showed dense fibers of GnRH3 in the pituitary (Fig. 2C-1), which should be able to regulate gonadotropin release as in other Characiform species that lost *gnrh1* [24] (e.g. all Characidae species examined in the present study), although some technical limitations prevented us from confirming the natural breeding ability of the *gnrh1* KO piranha.

### Both piranha gnrh1 and gnrh3 enhancers are active in the POA GnRH neurons expressing gnrh3 mRNA in zebrafish, although they do not have intrinsic gnrh1 gene

Given that piranha hypophysiotropic GnRH neurons co-express *gnrh1* and *gnrh3*, the common ancestor of Ostariophysi should be considered to have possessed hypophysiotropic neurons that co-express *gnrh1* and *gnrh3*. According to this hypothesis, the present ostariophysian fish that lack the *gnrh1* gene may be able to activate gene expression via the *gnrh1* enhancer of piranha in their hypophysiotropic GnRH neurons. By using zebrafish as a model, we demonstrated that both Tg (*pngnrh1*:RFP) and Tg (*pngnrh3*:GFP) zebrafish showed not only GFP (*gnrh3* enhancer induced) but also RFP (*gnrh1* enhancer induced) expression in the *gnrh3* mRNA-expressing hypophysiotropic neurons in the POA (Fig. 3), even though zebrafish has lost *gnrh1* gene probably in the common ancestor of Cyprinidae and Danionidae (75 Mya) [28] (Fig. 5). This result not only supports the hypothesis that *gnrh1* and *gnrh3* were co-expressed in hypophysiotropic GnRH neuron of their ancestor but also strongly suggests that transcription factors that can activate *gnrh1* enhancer have been conserved for an extended period of time in the absence of *gnrh1*. Given that *gnrh1* and *gnrh3* should have been transcribed by the same transcription factors at the time of duplication (1R/2R WGD), it is likely in zebrafish POA GnRH neurons that the transcription factors for intrinsic *gnrh3* could be activated by the piranha *gnrh1* enhancer. Although we could not specify the enhancer sequences because of low similarity in the 5’-flanking region sequence of *gnrh1* and *gnrh3*, a previous developmental study showing that the enhancers of paralogous genes was active in the same tissue [31] supports a hypothesis that *gnrh1* and *gnrh3* have possessed common cis- and trans-expression regulatory systems.

**Fig. 5.**
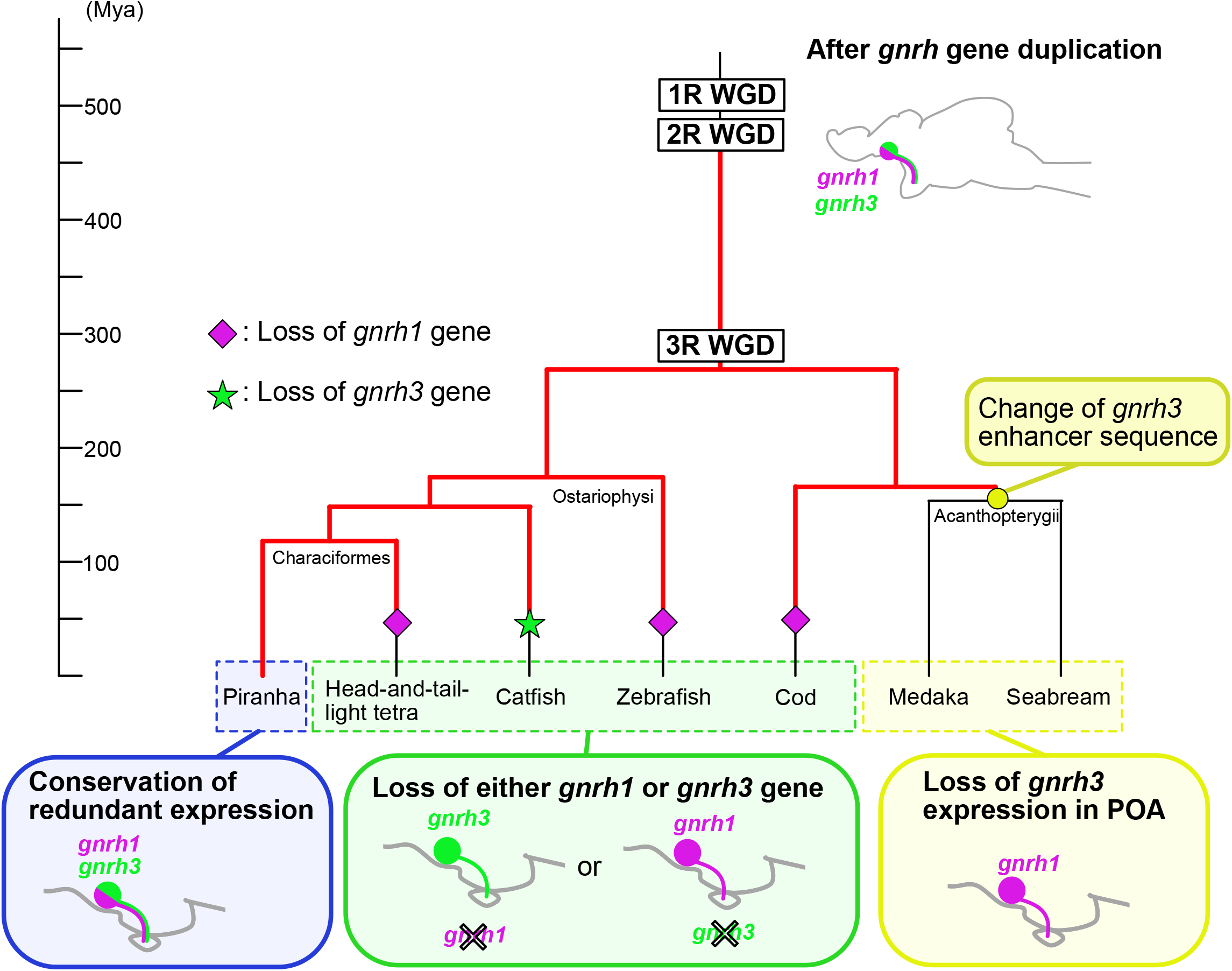
Working hypothesis of the evolution of paralogous *gnrh1/gnrh3* genes underlying the frequent switching of the *gnrh* gene expression in hypophysiotropic GnRH neurons. The present study provides evidence that piranha *gnrh1* and *gnrh3* are co-expressed in the hypophysiotropic GnRH neurons (blue box), which suggests that all its ancestors inherited the same property since the 1R/2R WGD. This evidence is the key to explaining why the loss of either *gnrh1* or *gnrh3* gene have been permitted in many ancestral teleosts. The red lines indicate hypothetical ancestors that co-expressed *gnrh1* and *gnrh3* in the hypophysiotropic GnRH neurons. Many other species so far examined in Ostariophysi lost either *gnrh3* or *gnrh1* (green box). Unlike other orders, in Acanthopterygii, the POA neuron-specific enhancer of *gnrh3* is suggested to have been lost in their ancestors (yellow circle), which is consistent with the experimental evidence that *gnrh1* is used in hypophysiotropic neurons in all species examined in Acanthopterygii (yellow box). Purple diamond and green star indicate loss of *gnrh1* and *gnrh3*, respectively.

### Both piranha gnrh1 and gnrh3 enhancers are active in the hypophysiotropic POA GnRH neurons of medaka expressing intrinsic gnrh1 mRNA

To uncover why most Acanthopterygii do not have hypophysiotropic *gnrh3* neurons in the POA despite possessing *gnrh3* gene itself [12, 13, 15, 17], we examined in medaka if their POA GnRH neurons are capable of activating gene expression via piranha *gnrh1* and *gnrh3* enhancers by generating transgenic medaka. Both Tg (*pngnrh1*:RFP) and Tg (*pngnrh3*:GFP) medaka showed enhancer activities of piranha *gnrh1* and *gnrh3* in POA *gnrh1* mRNA-expressing neurons (Fig. 4), although medaka does not express their intrinsic *gnrh3* mRNA in hypophysiotropic POA neurons [17]. The fact that piranha *gnrh3* enhancer was activated in *gnrh1* mRNA-expressing neurons in POA of medaka indicates that medaka GnRH1 neurons in the POA possess a transcription factor that is activated by the piranha *gnrh3* enhancer (Fig. 4C). In other words, the loss of POA-specific enhancer of *gnrh3* gene in a common ancestor can explain why only *gnrh1* is expressed in hypophysiotropic neurons in acanthopterygians including medaka. Interestingly, although all present-day species in Acanthopterygii possess both *gnrh1* and *gnrh3* [8, 20], *gnrh1* is exclusively expressed in the POA hypophysiotropic GnRH neurons in many species [12-17]. Especially, as *gnrh1* KO medaka leads to infertility due to the dysfunction in ovulation [32], *gnrh1* is exclusively important for the luteinizing hormone (LH) regulation [33]. Given the general situation in Acanthopterygii that *gnrh3* is seldom expressed in POA, the loss of the enhancer responsible for *gnrh3* expression in the POA neurons may have occurred in the ancestor of medaka in the early acanthopterygian lineage (∼150 Mya), which forced all acanthopterygian to use their remaining *gnrh1* in the hypophysiotropic GnRH neurons in the POA. On the other hand, non-acanthopterygian Euteleostei, Atlantic cod [34] have lost *gnrh1*, which indicate that the loss of the POA-specific enhancer of the *gnrh3* gene occurred at least after the divergence of Atlantic cod and Acanthopterygii (∼160 Mya) (Fig. 5).

In the present study, the enhancer activity of piranha *gnrh1* and *gnrh3* was analyzed in zebrafish/medaka TN neurons, in addition to the hypophysiotropic neurons. The enhancers of piranha *gnrh1* and *gnrh3* were also activated in *gnrh3* mRNA-expressing TN neurons in zebrafish (Fig. S4B, C). Also, the double transgenic zebrafish Tg (*pngnrh1*:RFP; *pngnrh3*:GFP) showed co-localization of RFP and GFP in some cells in the TN (Fig. S4D). It strongly suggests that zebrafish TN neurons have transcription factors to co-express *gnrh1* and *gnrh3* if they had an ancestral *gnrh1* gene. On the other hand, in medaka, reporter expression by the piranha *gnrh1* enhancer was not observed in TN *gnrh3* mRNA-expressing cells (Fig. S4E), while *pngnrh3*:GFP reporter expression was observed (Fig. S4F, G). In this case, we cannot distinguish whether these neurons do not have transcription factors to activate piranha *gnrh1* enhancer or an incompatibility of medaka transcription factors and piranha enhancer of this system prevents the transcription system.

On the other hand, all the other cases in the present study resulted in the demonstration of the piranha’s enhancer activity in medaka or zebrafish neurons. These clearly lead to a simple conclusion that the piranha *gnrh1/3* enhancer is active in zebrafish TN, POA, and medaka POA GnRH neurons, even though these species are phylogenetically distinct.

### The reason for prolonged conservation of redundant gnrh1 and gnrh3 expression after 1R/2R WGD

The present study revealed redundant co-expression of *gnrh1* and *gnrh3* in piranha hypophysiotropic POA GnRH neurons. Given that *gnrh1* and *gnrh3* arose in 1R/2R WGD, it implies that this co-expression has been inherited to every ancestor of piranha for ∼550 My. This long-lasting redundancy is surprising because redundant gene are generally eliminated immediately [3, 35].

According to simple gene dosage effects, increase in gene copies affects the amount of gene expression [36, 37]. In fact, adaptive increases in copy number have been reported in some genes during evolution [38, 39]. Similarly, in *gnrh* genes expressed in a hypophysiotropic neuron, this redundancy of *gnrh1* and *gnrh3* should be conserved under positive selective pressure, since the amount of the gene product directly affects the efficiency of ovulation and the number of offspring.

However, loss of *gnrh1* and *gnrh3* sometimes occurred in the teleost linage. These events suggest that the loss of either *gnrh1* or *gnrh3* was not critical for survival, although it may be weakly deleterious. According to a theory in population genetics, natural selection does not work theoretically when the population size is small [40], and weakly deleterious mutation may be fixed within a new population, also referred to as the founder effect [41]. For these reasons, loss of either the *gnrh1* or *gnrh3* gene/enhancer are considered to have occurred very slowly during the ∼550 My long history of vertebrate linage, and the extreme case may be piranha, which still conserves this redundancy even now (Fig. 5).

### Slowly progressing role-division of paralogous genes gnrh1 and gnrh3 provides a good model for understanding paralogous gene evolution

The findings of *gnrh* genes demonstrated in the present study give an intriguing example of the possible process of role-division of paralogous genes after duplication at the cellular level. Once genes are duplicated, they undergo neo-functionalization, sub-functionalization, or non-functionalization, leading their resulting paralogs to diverge into independent evolutionary paths, which usually prevents them from reuniting or swapping their roles. Unlike many genes that has undergone this process rapidly [3], *gnrh1*/*gnrh3* genes expressed in the hypophysiotropic GnRH neurons in vertebrates experienced this role-division process very slowly probably due to the weakly deleterious property of their loss, which resulted in the apparent switching of paralog usage during evolution. Also, GnRH may not be the only example of paralogous gene switching [42], and further cellular-level observation of paralogous genes across species may provide more information on the general rule of the speed of role-division during evolution. The present cellular-level study explains the mechanism of the evolutionary process of role-division of the very slowly evolving paralogous genes in hypophysiotropic neurons. These findings provide compelling evidence to support the genomics-based theories on paralogous genes and offer a deeper understanding of their evolution.

## Materials and Methods

### Animals

Piranha (red-bellied piranha, *Pygocentrus nattereri*) were obtained from local dealer or Suma Aqualife Park KOBE, and specimen that weighted more than 45 g were used for histological experiments. Male and female gonadosomatic indices (GSI) were 1.27 ± 0.90 and 6.33 ± 2.46, respectively. Head-and-tail-light tetra (*Hemigrammus ocellifer*, weight: 0.93 ± 0.37 g; GSI in male: 1.8, GSI in female: 13.7 ± 3.1) were obtained from a local dealer. RIKEN WT (RW) zebrafish (*Danio rerio*) and medaka (*Oryzias latipes*) were obtained from NBRP zebrafish and a local dealer, respectively, and sexually matured adult with weights > 270 mg (zebrafish) and > 110 mg (medaka) were used. All animals in this study were maintained at 14 h-light/ 10 h-dark at a water temperature of 27 ± 2 ºC. All experiments were conducted in accordance with the protocols approved by the Animal Care and Use Committee of the University of Tokyo (permission number, 17-1 and P19-3).

### Isolation of coding sequences and genomic sequences

Coding sequences of *gnrh1* and *gnrh3* genes of piranha were identified by 3’RACE with degenerate primers followed by 5’RACE of gene specific primers. Also, partial sequence of *gnrh3* of head-and-tail-light tetra was determined by 3’RACE followed by RT-PCR. The total RNA were isolated from piranha or head-and-tail-light tetra by using ISOGEN (Nippon gene, Tokyo, Japan), and were applied to the SMART RACE kit (Takara Bio USA, Mountain View, CA) according to the manufacturer’s instructions. After cloning into pGEM-T vector (Promega, Madison, WI), sequences were analyzed by a commercial Sanger sequence service. Degenerate primers of well-conserved coding regions that encode GnRH mature peptides. The primers used in these 5’ and 3’RACE are indicated in Table S1. Note that the degenerate primer that amplified piranha (Serrasalmidae) *gnrh1* could not amplify *gnrh1* of neon tetra, head-and-tail-light tetra or glowlight tetra (Characidae; Fig. S2H), which is consistent with the fact that a *gnrh1-*like gene is not found in a genome database of a Characidae fish, Mexican tetra. The isolation of the upstream genome sequences of *gnrh1* and *gnrh3* of piranha was performed using the Universal GenomeWalker ™ kit (Takara Bio USA) according to the instructions, and 5.0 kbp and 3.6 kbp upstream sequences were obtained, respectively.

### Preparation of the brain sections

Piranha, head-and-tail-light tetra, medaka and zebrafish were deeply anesthetized with 0.02% ethyl 3-aminobenzoate methanesulfonate (MS-222) (Sigma-Aldrich, St. Louis, MO) and fixed by perfusion with 4% paraformaldehyde (PFA) in PBS. For piranha and zebrafish, saline was perfused before the perfusion of the 4% PFA. After their brains were dissected out, they were post-fixed by 4% PFA in PBS for more than 4 hours. They were then immersed in 30% sucrose in PBS for more than 4 hours for cryoprotection. They were then embedded in 5% agarose (type IX-A; Sigma-Aldrich) and 20% sucrose in PBS, frozen in *n*-hexane (∼-60 °C), and serial frontal sections were prepared at 25 μm thick on a cryostat.

### Antibodies

Peptides of the precursor sites of piranha GnRH1 and GnRH3 (see Fig. S2A, B), called GAP sequence, conjugated to KLH were used as antigens, and rabbit polyclonal antibodies against them were produced in this study (outsourced to Sigma-Aldrich). The immunized species and dilution concentrations of the primary antibodies used are as follows; piranha GAP1 antibody (rabbit, 1/10,000) for piranha GnRH1; piranha GAP3 antibody (rabbit, 1/10,000) for piranha and head-and-tail-light tetra GnRH3; anti-RFP antibody 600-401-379S (rabbit, 1/2,000, Rockland Immunochemicals, Inc., Gilbertsville, PA) for RFP; GFP polyclonal antibody A11122 (rabbit, 1/1,000, Thermo Fisher Scientific, Waltham, MA) for GFP. For the secondary antibody, anti-rabbit IgG, biotinylated (goat, 1/200; Vector Laboratories, Burlingame, CA) was used.

### In situ hybridization, immunohistochemistry

For *in situ* hybridization, digoxigenin (DIG)-labeled RNA probes of piranha *gnrh1*, piranha *gnrh3*, head-and-tail-light tetra *gnrh3*, zebrafish *gnrh3*, medaka *gnrh1* and medaka *gnrh3* and fluorescein-labeled RNA probe of piranha *gnrh3* were synthesized with DIG or a fluorescein RNA labeling mix (Roche, Basel, Switzerland). After 1 μg/ml Proteinase K (Takara Bio, Shiga, Japan) treatment, the sections were hybridized with probes in a hybridization buffer (50% formamide, 3× SSC, 0.12 M phosphate buffer pH 7.4, 1× Denhardt solution, 125 μg/ml tRNA, 0.1 mg/ml calf thymus DNA, and 10% dextran sulfate) at 58 ºC overnight. After rinsing, for single-labeled *in situ* hybridization (piranha *gnrh1*, piranha *gnrh3*, head-and-tail-light tetra *gnrh3*), alkaline phosphatase-conjugated anti-DIG antibody (1/5,000, Roche) were treated and the alkaline phosphatase activity was detected by 337 μg/ml 4-nitroblue tetrazolium chloride (NBT) and 175 μg/ml 5-bromo-4-chloro-3-indoyl-phosphate (BCIP). For double-labeled *in situ* hybridization (piranha *gnrh1* and piranha *gnrh3*), alkaline phosphatase-conjugated anti-DIG antibody (1/2,000, Roche) and horseradish peroxidase (HRP)-conjugated anti-fluorescein antibody (1/500, Roche) were applied. The activity of alkaline phosphatase was visualized using SIGMAFAST™ Fast Red TR/Naphthol tablet (Sigma-Aldrich). HRP activity was detected by TSA-plus biotin (Pekin Elmer, Waltham, MA), followed by a Vectastain Elite ABC kit (Vector Laboratories), and then streptavidin, Alexa Fluor™ 488 conjugate (Thermo Fisher Scientific).

For immunohistochemistry of piranha GnRH1 and piranha and head-and-tail-light tetra GnRH3, sections were incubated in 0.5× SSC at 95 º C for 20 min for antigen retrieval before the primary antibody reaction. Primary antibody was incubated in PBS containing 5% normal goat serum at room temperature for immunohistochemistry of all samples (piranha GnRH1, piranha GnRH3 and head-and-tail-light tetra GnRH3). Afterwards, inactivation of endogenous peroxidase by 0.3% H_2_O_2_ and secondary antibody reaction were performed. The sections were incubated with Vectastain Elite ABC kit and the peroxidase activity was detected by 1 mg/ml 3,3-diaminobenzidine (DAB) and 0.003% H_2_O_2_. For validation of produced piranha GAP1 and GAP3 antibodies, immunohistochemistry was performed using preabsorbed primary antibody with 0.01% KLH and 1 μM GAP1 or GAP3 peptide.

By using transgenic zebrafish and medaka brain sections, double labeling of *in situ* hybridization and immunohistochemistry was performed. Primary antibody against RFP or GFP was treated at 4ºC overnight, and for immunohistochemistry of RFP, 1 μg/ml Proteinase K were treated for the antigen retrieval of RFP epitope before primary antibody reaction. After secondary antibody reaction, samples were fixed with 4% PFA, and hybridization with a DIG RNA probe (*gnrh1* or *gnrh3* of zebrafish or medaka) was performed in a hybridization buffer described above at 58 ºC overnight. After rinsing, peroxidase-conjugated anti-DIG antibody (0.3 U/ml, Roche) were treated, and the peroxidase activity was detected by TSA Plus Cyanine 3 kit (Perkin Elmer) or Alexa Fluor™ 488 Tyramide SuperBoost™ Kit (Thermo Fisher Scientific). Afterwards, the signal of the GFP or RFP was visualized by Vectastain Elite ABC kit followed by streptavidin-conjugated Alexa Fluor™ 488 or Alexa Fluor™ 555 (Thermo Fisher Scientific).

For dual immunohistochemistry, brain sections of transgenic zebrafish and medaka were treated with Proteinase K for antigen treatment (37 ºC, 10 min). They were then incubated with primary antibody against RFP raised in mouse (1/2,000; RF5R, Thermo Fisher Scientific) and antibody against GFP raised in rabbit (1/1,000; A11122, Thermo Fisher Scientific) at room temperature overnight. After secondary antibody treatment (Alexa Fluor™ 555 conjugated anti-mouse IgG, (1/800; Thermo Fisher Scientific) and anti-rabbit IgG, biotinylated (1/200; Vector Laboratories) for 2 hours, the sections were incubated with Vectastain Elite ABC kit followed by streptavidin, Alexa Fluor™ 488 conjugate. Fluorescence-labeled sections were mounted on coverslips with CC/Mount (Sigma-Aldrich). Sections with DAB labeling were captured by an upright microscope, Olympus BX-53 equipped with Wraymer NOA630B or NOA2000. All the fluorescence-labeled sections were observed under a confocal microscope, Olympus FV1000. Contrasts and brightness were modified with ImageJ (NIH, Bethesda, MD).

### Artificial fertilization and generation of gnrh1 KO piranha

Mature female piranha were given intraperitoneal injection of 3 or 5 U/g body weight human chorionic gonadotropin (hCG, ASKA Animal Health, Tokyo, Japan) twice every 24 hours. Unfertilized eggs were obtained by squeezing the abdomen 12 hours after the second injection. After anesthesia with MS-222, a portion of the testis was removed through an incision in the abdomen of the male and a testicular suspension was prepared. Fertilized eggs were obtained by quickly mixing the unfertilized eggs and the testicular suspension. To generate KO piranha using the CRISPR/Cas9 system, we designed three types of gRNA at the 5’ side of the sequence of the *gnrh1* mature peptide. A mixture containing 30 ng/μl of each gRNA, 1 μg/μl of Cas9 protein, 100 ng/μl of tracr RNA, 1 ng/μl of EGFP mRNA, and 0.02% phenol red in 1× PBS was injected into piranha fertilized eggs. The designed gRNA regions are shown in Fig. S2A. The primers for genotyping are shown in Table S1.

### Generation of transgenic zebrafish and medaka

The upstream sequences of cloned piranha *gnrh1* or *gnrh3* were connected to zebrafish heat shock promoter as a minimal promoter, dTomato (RFP; for piranha *gnrh1*) or EGFP (GFP; for piranha *gnrh3*) and bovine growth hormone poly(A) signal sequentially and inserted into the pGEM-T vector. For screening in embryos, cardiac myosin light chain 2 promoter of zebrafish, mCherry (RFP; for piranha *gnrh1*) or EGFP (for piranha *gnrh3*) and SV40 poly(A) signal were inserted downstream of reporter construct [43]. Microinjection of the plasmid into fertilized eggs of zebrafish and medaka was performed. After maturation, they were crossed and F1 embryos were screened based on the fluorescence in cardiac muscle.

## Acknowledgments

We thank Nobuyuki Higashiguchi and Nobuhito Osada (Suma Aqualife Park KOBE) for providing red-bellied piranha. We are also grateful to Dr. Shin-ichi Higashijima (National Institute for Basic Biology, Japan) for providing plasmid containing zebrafish heat shock promoter sequence. We are indebted to Dr. Yukinori Kazeto (National Research Institute of Aquaculture, Fisheries Research and Education Agency, Japan) for helpful advice in the induction of ovulation in teleosts. We also thank Drs. Tatsuo Michiue and Yuko Mochizuki (the University of Tokyo) for helpful advice in artificial insemination. We are also grateful to Drs. Mikoto Nakajo (Osaka Medical and Pharmaceutical University) and Chie Umatani (Tokyo University of Agriculture and Technology) for helpful discussion and comments.

## Funding

Research grants in the Natural Sciences of Mitsubishi Foundation (SK)

Grant for Basic Science Research Projects of Sumitomo Foundation (SK)

Japan Society for the Promotion of Science 18K19323 (SK)

Japan Society for the Promotion of Science 18H04881 (SK)

Japan Society for the Promotion of Science 24657050 (YO)

## Author contributions

Conceptualization: CF, SK

Investigation: CF, KoS, MI, CY, DK, ST, KaS, YA, SK

Supervision: SK

Writing—original draft: CF, SK

Writing—review & editing: CF, CY, DK, ST, YA, YO

Project administration: SK

Funding acquisition: YO, SK

## Data and materials availability

All data are available in the main text or the supplementary materials.

