## Supplementary material for "Enhancer evolution as a driving force for lineage-specific paralog usage in the central nervous system": Fig. S

### Supplemental figure1

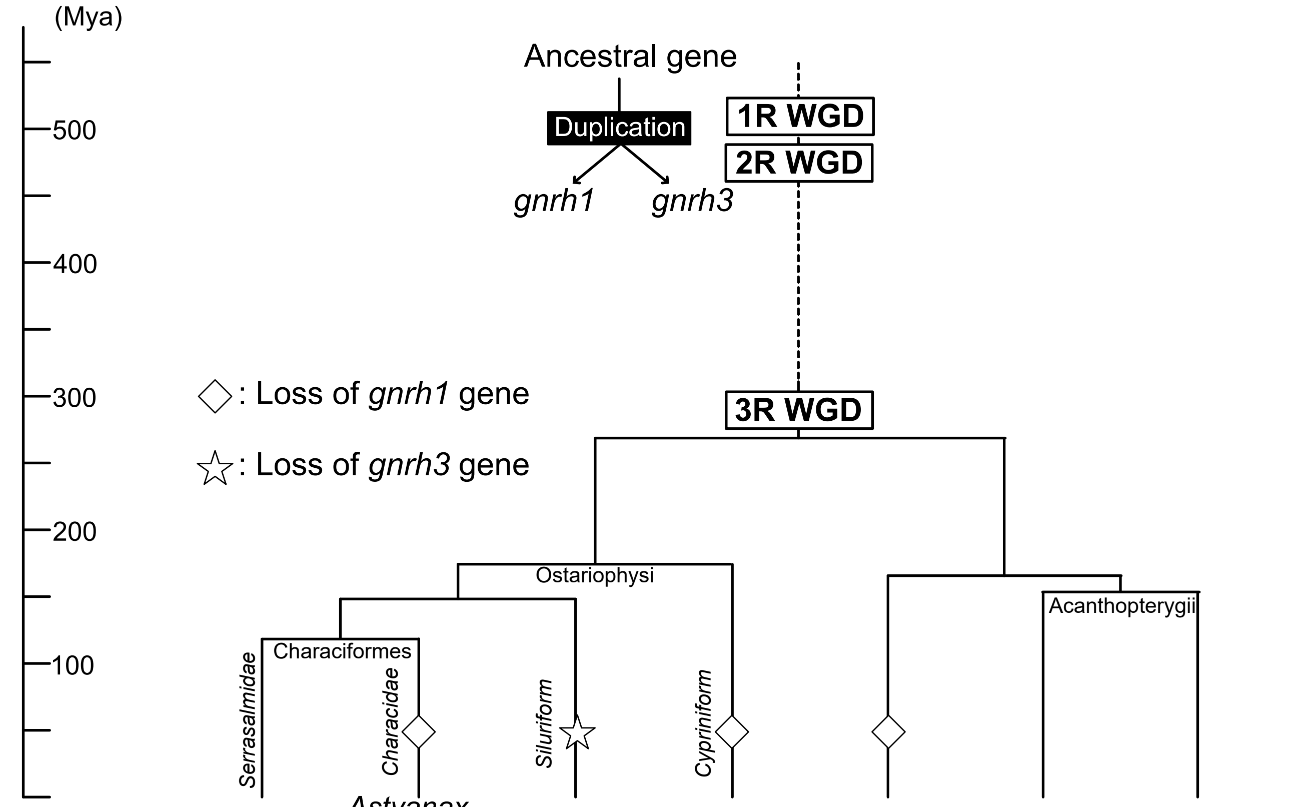


| | Piranha | *Astyanax altiparanae* | Catfish | Zebrafish | Cod | Medaka | Seabream |
|---|---|---|---|---|---|---|---|
| *gnrh* paralog | *gnrh1*<br>*gnrh3* | *gnrh3* | *gnrh1* | *gnrh3* | *gnrh3* | *gnrh1*<br>*gnrh3* | *gnrh1*<br>*gnrh3* |
| (reference) | (this study) | (1) | (2) | (4) | (6) | (7) | (9) |

| | Piranha | *Astyanax altiparanae* | Catfish | Zebrafish | Cod | Medaka | Seabream |
|---|---|---|---|---|---|---|---|
| Hypophysiotropic *gnrh* | ? | *gnrh3 ?* | *gnrh1* | *gnrh3* | *gnrh3?* | *gnrh1* | *gnrh1* |
| (reference) | | | (3) | (5) | | (8) | (10) |

**Fig. S1. Phylogenetic tree of teleost species and hypophysiotropic GnRH paralogs.**
The *gnrh1* and *gnrh3* genes were generated from an ancestral gene during the first/second-round (1R/2R) whole genome duplication (WGD), which occurred ~550 million years ago (Mya). Many teleost species have hypophysiotropic GnRH1 neurons as the main regulator of gonadotropin release in Acanthopterygii. However, in Cypriniform and Characidae species as well as Atlantic cod, *gnrh1* has been lost and *gnrh3* is expressed in hypophysiotropic GnRH neurons instead. Since three forms of GnRH peptide are expressed in a Serrasalmidae species, pacu, we expected species in this group, including piranha, may have both *gnrh1* and *gnrh3*.

### Supplemental figure 2

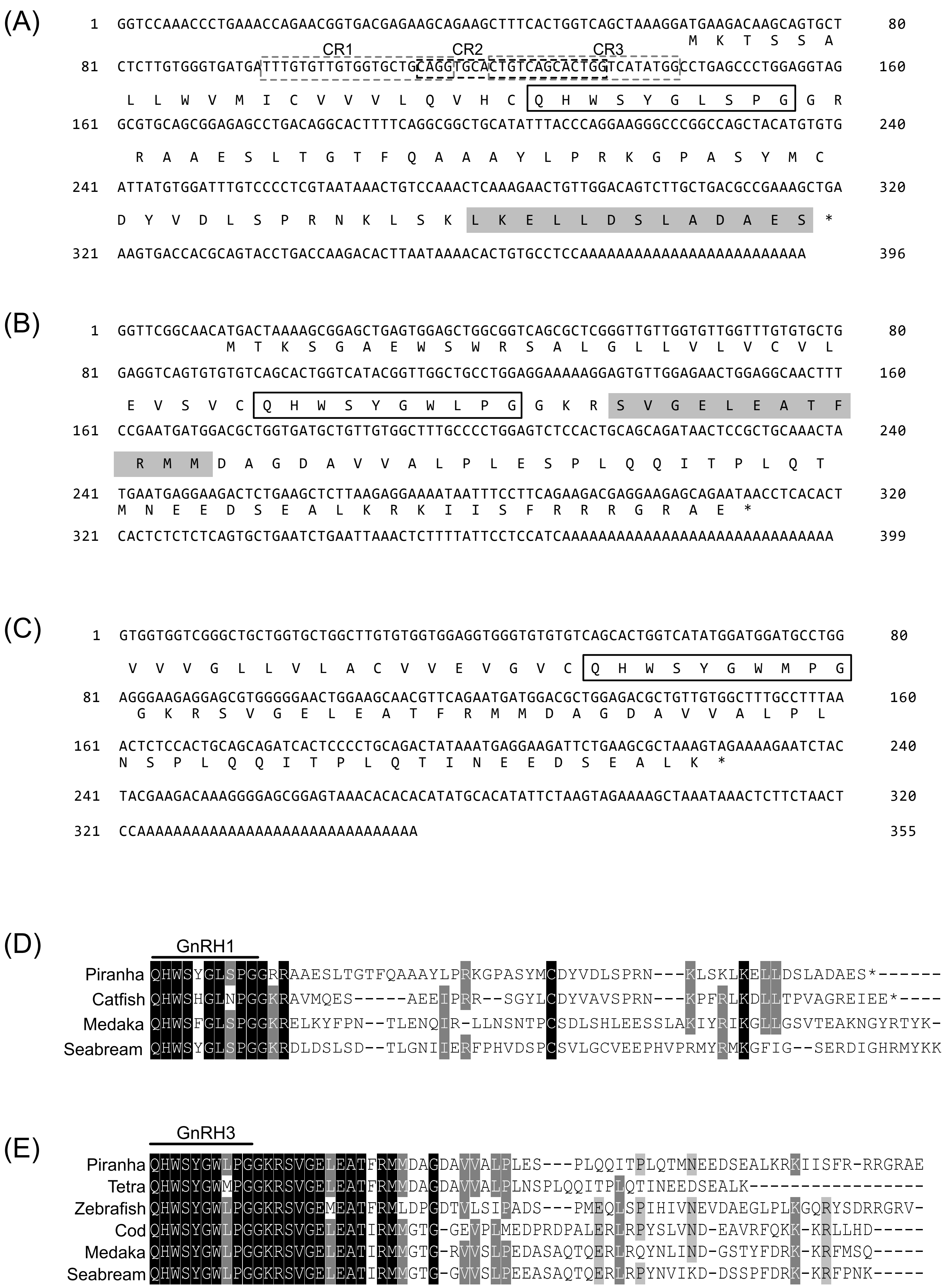

(F)

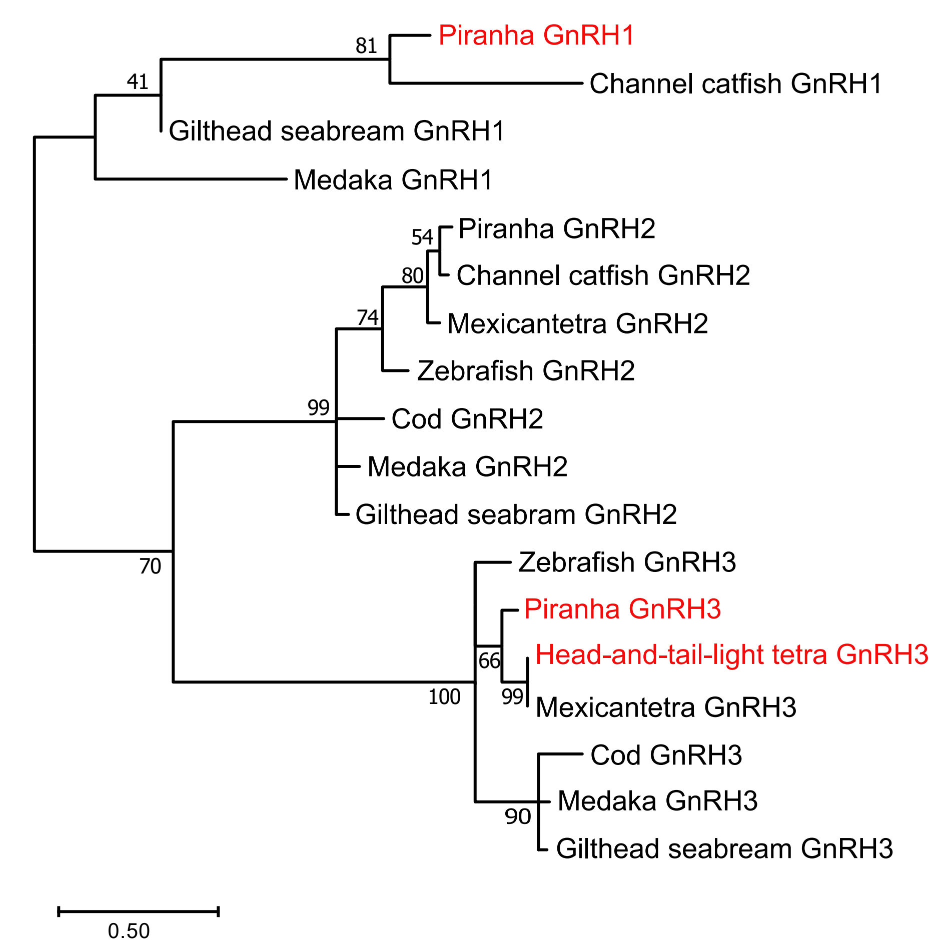


(G)

```
Wild type      TGTCAGCACTGGTCATATGGCCTGAGCCCTGGA
                    Q  H  W  S  Y  G  L  S  P  G

1bp deletion   TGTCAGCACTGGT-ATATGGCCTGAGCCCTGGA
                    Q  H  W    Y  M  A  *

10bp deletion  TGTCA----------TATGGCCTGAGCCCTGGA
                              H  M  A  *

9bp deletion   TG---------GTCATATGGCCTGAGCCCTGGA
                          W  S  Y  G  L  S  P  G
```

(H)

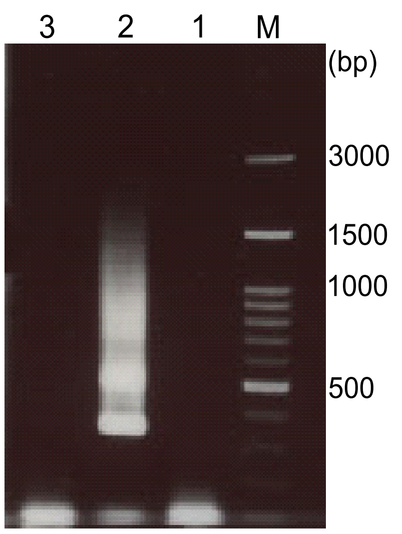

**Fig. S2. Alignment and phylogenetic tree of piranha *gnrh1*, *gnrh3* and head-and-tail-light tetra *gnrh3*.**

(**A**) Nucleotide sequence and deduced amino acid sequence of piranha *gnrh1* gene. Box shows mature GnRH peptides. Three dotted boxes (CR1, CR2, CR3) represent gRNA sequences used for generation of *gnrh1* knockout (KO) piranha. Antigen sequence for antibody production is shaded in gray. (**B**) Nucleotide sequence and deduced amino acid sequence of piranha *gnrh3* gene. Box shows mature GnRH peptides. Antigen sequence for antibody production is shaded in gray. (**C**) Partial nucleotide sequence and its deduced amino acid sequence of head-and-tail-light tetra *gnrh3* gene. Box shows mature GnRH peptides. (**D**) Alignment of the deduced amino acid sequence of mature GnRH1 and GnRH-associated peptide (GAP1) in piranha with those in channel catfish (catfish), medaka and gilthead seabream (seabream). Mature GnRH1 sequence of piranha is identical to that of gilthead seabream. Conserved amino acids in all 4 species and 3 species are shaded black and gray, respectively. (**E**) Alignment of the deduced amino acid sequence of mature GnRH3 and GnRH-associated peptide (GAP3) in piranha and head-and-tail-light tetra with those in zebrafish, cod, medaka and gilthead seabream (seabream). Mature GnRH3 sequence of piranha is identical to that of all species aligned except for head-and-tail-light tetra (Tetra). Conserved amino acids in all 6 species and 4 species are shaded black and gray, respectively. (**F**) Phylogenetic tree of GnRH paralogs in piranha, head-and-tail-light tetra, and other teleosts (Maximum likelihood). The number at each node shows the percentage of the bootstrap value for 1000 replicates. Scale bar represents 0.50 substitutions per site. (**G**) Genomic sequence and deduced mature peptide sequence of wild type and KO *gnrh1* gene. Both 1-bp and 10-bp deletion mutation resulted in a frameshift and a premature stop codon (asterisk). In all mutations, it is expected that functional GnRH peptides were not produced. **(H)** Detection of *gnrh1* mRNA was failed in various Characidae fish cDNA. Degenerate primers that amplified piranha *gnrh1* failed to amplify *gnrh1* of neontetra (lane1), head-and-tail-light tetra (lane2) and glowlight tetra (lane3). Note that the bands observed in lane 2 are nonspecific amplicons. M, DNA markers (bp).

### Supplemental figure 3

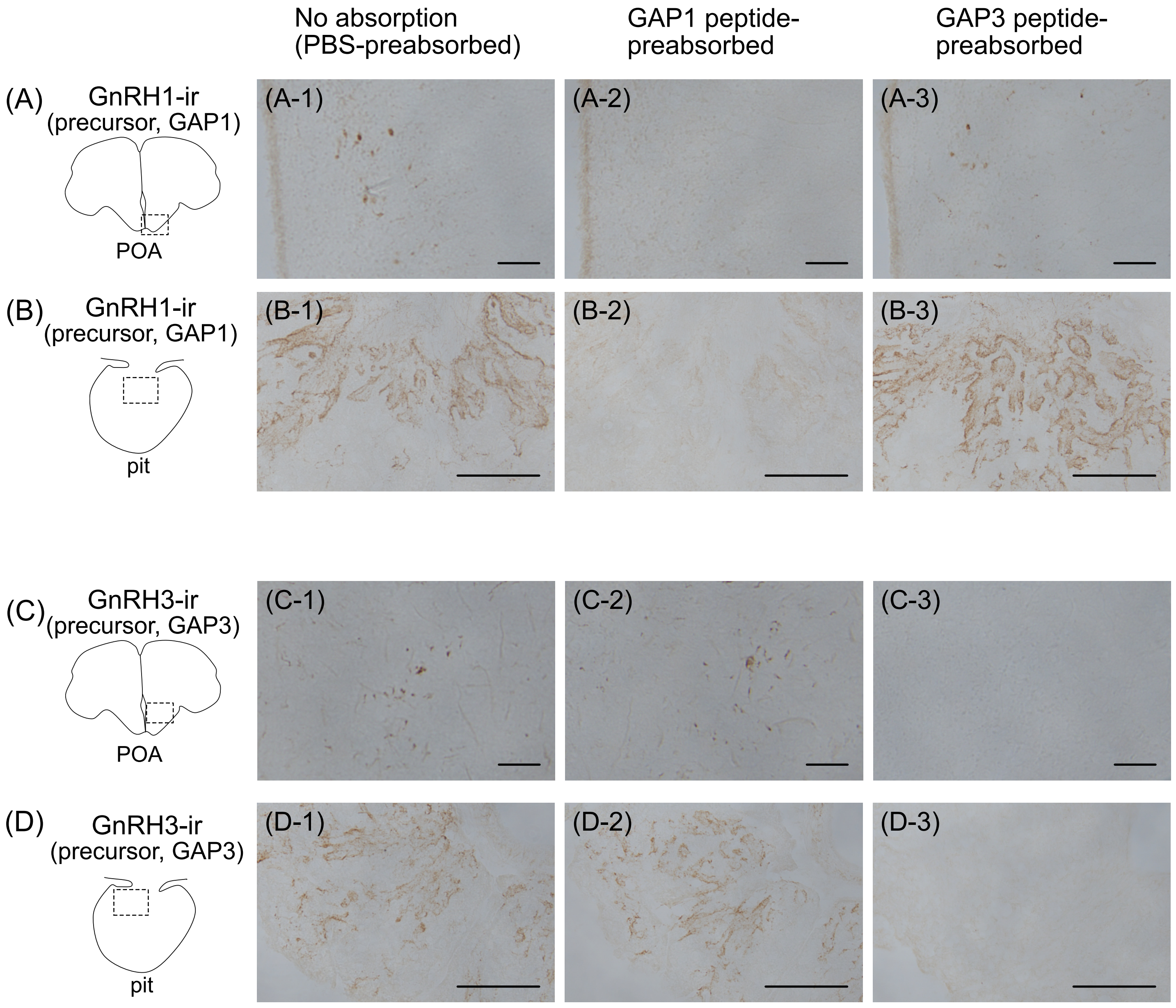

**Fig. S3. Specificities of produced GnRH precursor antibodies.**

(**A, B**) Immunohistochemistry for GnRH1 precursor of transverse section of piranha POA (**A**) and pituitary (**B**). Immunoreactive cells and fibers are labeled using a GnRH1 precursor antibody (**A-1, B-1**) and are not labeled with a GnRH1 precursor antibody preabsorbed with GnRH1 precursor peptides (**A-2, B-2**). (**A-3, B-3**) In immunohistochemistry using a GnRH1 precursor antibody preabsorbed with GnRH3 precursor peptides, immunoreactive cells and fibers are labeled. (**C, D**) Immunohistochemistry for GnRH3 precursors showed that immunoreactive cells and fibers are labeled in the POA (**C**) and pituitary (**D**) when preabsorbed with PBS (**C-1, D-1**) or GnRH1 precursor peptides (**C-2, D-2**), but are not labeled when preabsorbed with GnRH3 precursor peptides (**C-3, D-3**). Scale bars, 100 μm.

### Supplemental figure 4

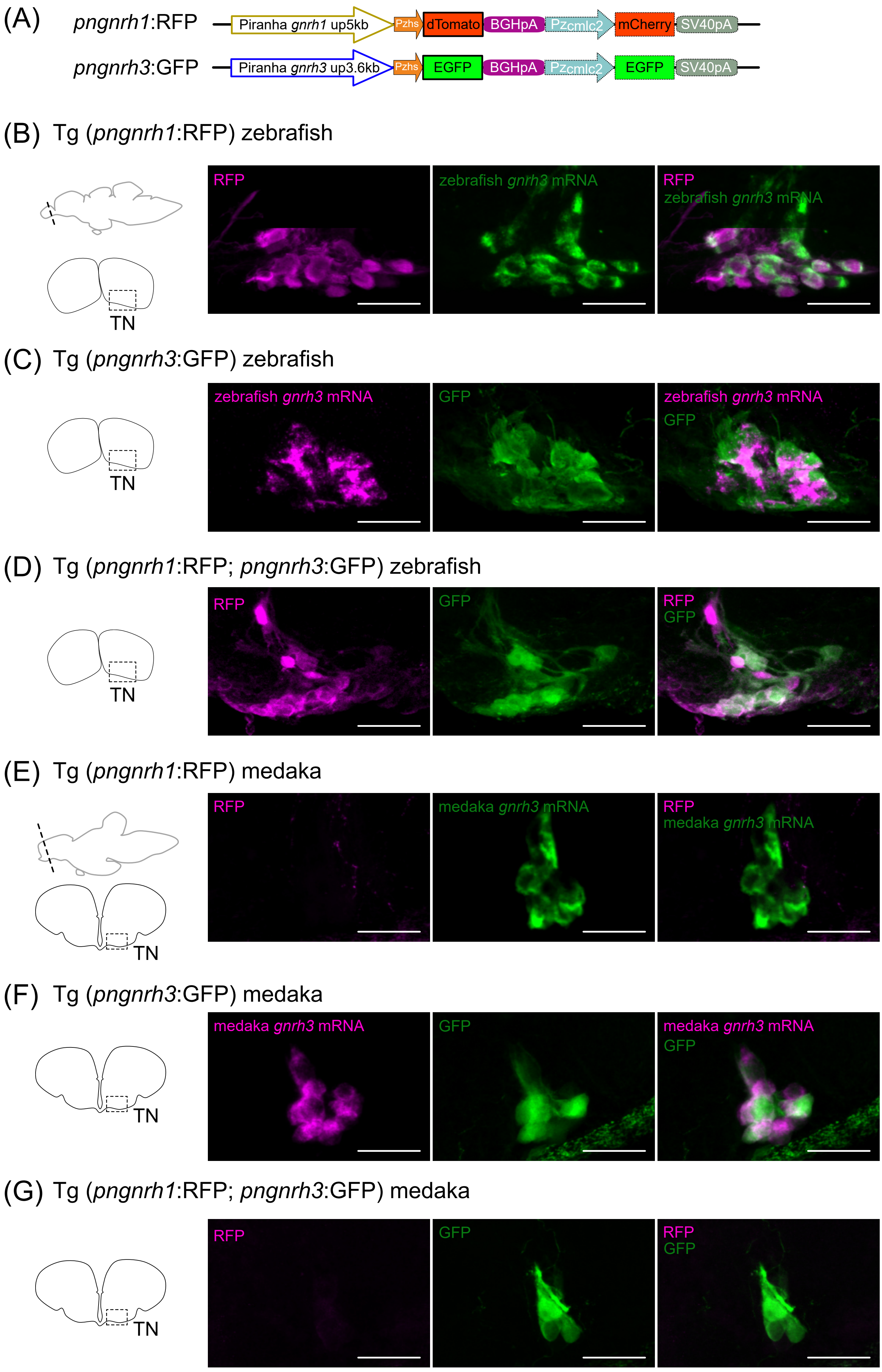

**Fig. S4. Examination of enhancer activity of piranha *gnrh1* and *gnrh3* (*pngnrh1*/*pngnrh3*) in the terminal nerve (TN) of zebrafish and medaka.**

(**A**) The constructs used to generate transgenic zebrafish and medaka. Both constructs examine the enhancer activity of piranha *gnrh1* or *gnrh3* 5'flanking regions by using a basal promoter (zebrafish heat shock promoter, Pzhs) and a fluorescent protein (dTomato or EGFP). For screening of embryos, cardiac myosin light chain 2 promoter of zebrafish (Pzcmlc2), mCherry or EGFP and SV40 poly(A) signal were inserted downstream of reporter construct. (**B**) In Tg (*pngnrh1*:RFP) zebrafish, *pngnrh1* enhancer-induced RFP expression is observed in the GnRH3 neurons (*gnrh3* mRNA-expressing neurons) in the TN. (**C**) In Tg (*pngnrh3*:GFP) zebrafish, *pngnrh3* enhancer-induced GFP expression is observed in the GnRH3 neurons in the TN. (**D**) Analysis of the double transgenic zebrafish, Tg (*pngnrh1*:RFP; *pngnrh3*:GFP). In Tg (*pngnrh1*:RFP; *pngnrh3*:GFP) zebrafish, some of the neurons in the TN express both RFP and GFP suggesting that *pngnrh1* and *pngnrh3* enhancers are activated in the same neurons. (**E**) In Tg (*pngnrh1*:RFP) medaka, *pngnrh1* enhancer-induced RFP expression is not observed in the GnRH3 neurons (*gnrh3* mRNA-expressing neurons) in the TN. (**F**) In Tg (*pngnrh3*:GFP) medaka, *pngnrh3* enhancer-induced GFP expression is observed in the GnRH3 neurons in the TN. (**G**) In double transgenic medaka, Tg (*pngnrh1*:RFP; *pngnrh3*:GFP), some of the neurons in the TN express only GFP suggesting that only *pngnrh3* enhancers are activated in the TN. Scale bars, 100 μm.

**Table S1. The sequence of primers used in this study.**

For full-length cloning of piranha *gnrh1*

5'-CTSYCAGCAYTGGTCITWYGG-3'
5'-ACTGGTCNTWYGGICTIMGICCIGGIG-3'
5'-TACTGCGTGGTCACTTTCAGCTTTCGG-3'
5'-TGCAGCCGCCTGAAAAGTGCCTGTC-3'

For full-length cloning of piranha *gnrh3*

5'-ACCCTSTSYCARCAYTGGTCITAYGGITGG-3'
5'-AGCACTGGTCIYAYGGITGGYWNCCIGGIGG-3'
5'-CARCAYTGGTCITAYGGITGG-3'
5'-TATCTGCTGCAGTGGAGACTCCAGG-3'
5'-GGGGCAAAGCCACAACAGCATCACC-3'

For cloning of head-and-tail-light tetra *gnrh3*

5'-CTSYCAGCAYTGGTCITWYGG-3'
5'-ACTGGTCNTWYGGICTIMGICCIGGIG-3'
5'-CTACTTTAGCGCTTCAGAATCTTCCTCA-3'
5'-TCTCCAGCGTCCATCATTCTGAA-3'
5'-GTGATCTGCTGCAGTGGAGAGTT-3'

For genotyping of piranha *gnrh1* KO

5'-GGCTGTTTTCTGTTGCTCTTTTCAGGATG-3'
5'-TACCGCCTGAAAAGTGCCTGTCA-3'
